# SARS-CoV2 envelop proteins reshape the serological responses of COVID-19 patients

**DOI:** 10.1101/2021.02.15.431237

**Authors:** Sophie Martin, Christopher Heslan, Gwénaële Jégou, Leif A. Eriksson, Matthieu Le Gallo, Vincent Thibault, Eric Chevet, Florence Godey, Tony Avril

## Abstract

The SARS-CoV-2 pandemic has elicited a unique international mobilization of the scientific community to better understand this coronavirus and its associated disease and to develop efficient tools to combat infection. Similar to other *coronavirae*, SARS-CoV-2 hijacks the host cell complex secretory machinery to produce properly folded viral proteins that will compose the nascent virions; including Spike, Envelope and Membrane proteins, the most exposed membrane viral proteins to the host immune system. Antibody response is part of the anti-viral immune arsenal that infected patients develop to fight viral particles in the body. Herein, we investigate the immunogenic potential of Spike (S), Envelope (E) and Membrane (M) proteins using a human cell-based system to mimic membrane insertion and N-glycosylation. We show that both S and M proteins elicit the production of specific IgG, IgM and IgA in SARS-CoV-2 infected patients. Elevated Ig responses were observed in COVID+ patients with moderate and severe forms of the disease. Finally, when SARS-CoV-2 Spike D614 and G614 variants were compared, reduced Ig binding was observed with the Spike G614 variant. Altogether, this study underlines the needs for including topological features in envelop proteins to better characterize the serological status of COVID+ patients, points towards an unexpected immune response against the M protein and shows that our assay could represent a powerful tool to test humoral responses against actively evolving SARS-CoV-2 variants and vaccine effectiveness.

## Introduction

The current COVID-19 pandemic has triggered unprecedented collective research efforts from the scientific community to better understand the disease and its cellular and molecular mechanisms, to identify efficient therapeutic drugs for taking care of infected patients with the severe acute respiratory syndrome coronavirus 2 (SARS-CoV-2), and to develop vaccines for protecting the whole population from the infection (1–3). One of the initial challenges in the fight against this virus was to rapidly detect SARS-CoV-2 infected patients to limit the propagation of the virus through isolation (4). Another challenge was to better understand the global antibody responses against SARS-CoV-2 proteins in patients (2, 5).

Among the anti-viral immune responses elicited in infected patients, immunoglobulin (Ig) responses against viral membrane proteins expressed at the surface of the virus are important for generating antibodies that limit virus propagation. This occurs by preventing interactions with host cells, *i.e.* production of neutralizing antibodies that block the binding of the viral membrane Spike protein to its receptor angiotensin-converting enzyme 2 (ACE2) expressed by infected host cells (2, 5–7). These anti-virus antibodies are also key mediators to trigger antibody-dependent immune responses such as the complement-dependent cytotoxicity as part of the humoral response (5) or the antibody-dependent cellular cytotoxicity mediated by immune cells harboring Fc receptors such as NK lymphocytes, macrophages and granulocytes to allow phagocytosis and destruction of the virus (5).

Some of these cellular actors such as macrophages and neutrophils could also contribute to the aggravation of the disease by releasing chemokines and cytokines that enhance inflammatory cascades described as ‘cytokine storms’ leading to lesions of infected tissues; although the involvement of the antibody-dependent mechanisms still need to be confirmed in COVID-19 patients (5). Most of the serological assays developed against SARS-CoV-2 are based on the recognition of the viral membrane Spike molecule and the nucleocapsid protein N, considered as major targets of antibody responses (8, 9). Besides the viral Spike molecule, little is described on Ig responses against the others viral membrane proteins E and M also directly exposed to the host immune system.

Spike (S), envelope (E) and membrane (M) are integral membrane proteins that transit through the host cells’ endoplasmic reticulum (ER). In this compartment, they are N-glycosylated, folded, and assembled in the ER-Golgi Intermediate compartment for virus budding and release (3). This maturation process is key for proper viral protein functions. For instance, Spike N-linked glycosylation is required for virus entry into the host cells impacting directly on Spike stability during its synthesis instead of its binding ability to the ACE2 receptor (10). These modifications might be also key for antibody recognition.

In the present study, we relied on an experimental system that recapitulates protein modifications acquired through the host cells’ secretory pathway to explore the antibody responses of SARS-CoV-2 infected patients. We found that S and M proteins (but not E) exhibited antigenic domains recognized by IgG, IgM and IgA in SARS-CoV-2 infected patients. High levels of Ig responses were observed in COVID-19 patients with moderate and severe forms of the disease. Finally, SARS-CoV-2 Spike D614 and G614 variants were compared, showing reduced Ig binding on the Spike G614 variant. Altogether, this study underlines the necessity of considering the mammalian cellular system to better characterize the serological status of COVID-19 patients.

## Materials and Methods

### Antibodies, plasmids and other reagents

All antibodies except those specified below were purchased from Jackson Immunoresearch (Ozyme, Saint-Cyr-L’École, France). We also used the rabbit monoclonal anti-SARS-CoV-2 Spike S1 (Sino Biologicals, Clinisciences, Nanterre, France) antibody. The following secondary antibodies were used: Alexa Fluor (AF) 488 conjugated donkey anti-rabbit IgG, AF488 conjugated F(ab’)2 donkey anti-human IgG, AF647 F(ab’)2 donkey anti-human IgM, FITC conjugated F(ab’)2 goat anti-human IgA (Thermo Fisher Scientific, Illkirch, France), Brilliant Violet 650 conjugated streptavidin (BioLegend, Ozyme), horseradish peroxidase (HRP) conjugated polyclonal goat anti-rabbit IgG (Dako, Agilent, Les Ulis, France), HRP conjugated StrepTactin (IBA GmbH, Fisher Scientific, Illkirch, France), and HRP-conjugated anti-FLAG (Sigma-Aldrich (St Quentin Fallavier, France)). Plasmids pTwist EF1alpha nCoV-2019 S 2xStrep, pLVX EF1alpha nCoV2019 E IRES-Puro and pLVX EF1alpha nCoV2019 M IRES-Puro encoding for SARS-CoV-2 Spike (D614 variant), E and M proteins respectively were a kind gift from the Krogan laboratory (UCSF, San Francisco, CA, USA) (11); and pCMV3 nCoV2019 Spike (D614 variant) C-FLAG Hygro was obtained from Addgene (Teddington, UK). Plasmid encoding for Spike G614 variant was generated using the pTwist EF1alpha nCoV-2019 S 2xStrep plasmid and the Q5 site-directed mutagenesis kit (New England BioLabs, Evry, France) following the manufacturer’s recommendations. D614 (codon GAC at position 1849) was replaced by G614 (codon GGC at the same position) with the following primers (IDT, Leuven, Belgium): forward 5’-CTTTATCAGGgCGTGAATTGTAC-3’ and reverse 5’-AACTGCAACCTGATTACTG-3’. The sequence of the modified plasmid was further verified after complete sequencing (Integragen, Evry, France). Other reagents not specified below were purchased from Sigma-Aldrich.

### Human sera collection

The study was carried out according to the regulation of Rennes Biobank (BRIF number: BB-0033-00056) certified as meeting the requirements of NF S96900 for receipt preparation preservation and provision of biological resources. Serum samples were gathered in the SEROCOV collection (DC-2019-3585). Socio-demographic information, underlying medical conditions, history of symptoms back to January 2020, and history of COVID-19 diagnosis before this investigation were collected at the time of the blood test and were presented in **Tables 1** and **S1**. Each COVID-19 participant was documented by a positive SARS-CoV-2 RT-PCR on respiratory samples. COVID-19 patients were categorized according to their symptom’s status based on their clinical conditions and care requirement. Patients with symptoms (fever, cough, anosmia, dysgeusia,…) and who did not require hospitalization were classified as mild COVID-19. Patients with symptoms and requiring hospitalization for oxygen therapy were classified as moderate COVID-19. Main patients of this group were cared for in pneumology, emergency (ENT), polyvalent internal medicine and geriatric units. Severe forms of COVID-19 were defined by patients requiring intensive care unit (ICU) hospitalization and oxygen therapy (oxygen flow superior to 6L/min or intubated). Patients with hyper-immunoglobulin M syndromes presented lupus pathology with cryoglobulinemia or primary parvovirus B19/EBV infection. Five sera were selected from infected patients with classical seasonal coronaviruses including 3/5 hCoV-OC43, 1/5 hCoV-NL63 and 1/5 hCoV-229E. Pre-pandemic sera were collected residual samples drawn before January 2020; and SARS-CoV-2 infected patients heparinized plasma were obtained from hospitalized patients at Rennes University Hospital Pontchaillou and the Centre Eugène Marquis (Rennes, France) between March 11^th^ and September 15^th^, 2020. All sera were aliquoted and conserved at 4°C for short-term use or frozen at −80°C.

**Table 1.** Clinical features of the cohort used in this study.

|  | **Age**<br>(mean±SD) | **Sex**<br>(% F / M) | **Time from**<br>**PCR** | **Time from**<br>**symptoms** | **Symptoms**<br>**/ treatments** |
| --- | --- | --- | --- | --- | --- |
| **before 2020/01** |  |  |  |  |  |
| **CTR** (n=38) | 37±14.8 | 54 / 46 | - | - | - |
| **CTR2** | 33±28.8 | 50 / 50 | - | - | - |
| coronavirus<br>(n=5) | 45±25.1 | 25 / 75 | 59±39.3 | - | - |
| hyperIg M<br>syndrome<br>(n=5) | 22±30.5 | 75 / 25 | - | - | - |
| **after 2020/01** |  |  |  |  |  |
| no symptom<br>(n=26) | 38±10.4 | 62 / 38 | - | - | 2 donors were<br>contact persons |
| symptoms<br>(n=4) | 50±18.9 | 50 / 50 | 130±21.6 |  | cough, fatigue, fever |
| **COVID+** |  |  |  |  |  |
| mild<br>(n=22) | 48±26.8 | 77 / 23 | 41±53.2 | 52±63.6 | cough, fatigue,<br>fever, anosmia |
| moderate<br>(n=14) | 72±16.1 | 42 / 58 | 14±14.7 | 25±16.8 | lung damages<br>/ hospitalization,<br>O<sub>2</sub> therapy (2.6±0.96) |
| severe<br>(n=17) | 64±11.2 | 34 / 66 | 15±8.1 | 25±8.2 | - / hospitalisation,<br>O<sub>2</sub> therapy (11±6.2),<br>intubated |

### Cell culture and transfection

Human epithelial HEK293T (HEK) cells were grown in Dulbecco’s modified Eagle’s medium (Gibco, Thermo Fisher Scientific) supplemented with 10% heat-inactivated fetal bovine serum (FBS) in a 5% CO2 humidified atmosphere at 37°C. For transient overexpression of SARS-CoV-2 membrane proteins, HEK cells (10^6^) were seeded in a 10 cm Petri dish with 10 mL complete medium for 24 hours, and then transfected using calcium phosphate co-precipitation with DNA for 48 hours. Plasmids (10 μg per dish) were initially diluted with 0.5 mL of CaCl2 (120 mM) and 0.5 mL of HEPES Buffer Saline solution (2x: HEPES 55 mM, NaCl 274 mM, Na2HPO4 1.4 mM, pH 7.05).

### Western blotting

SARS-CoV-2 S, E and M expressing HEK cells were resuspended in ice-cold lysis buffer (composed of 20 mM Tris-HCl, pH 7.5, 150 mM NaCl, 1% Triton X-100) supplemented with protease and phosphatase inhibitor cocktails (Roche, Sigma-Aldrich). Proteins were resolved by SDS-polyacrylamide gel electrophoresis (12% and 7% polyacrylamide gels for viral E and M proteins, and S proteins respectively) and transferred to nitrocellulose membrane for blotting. The membranes were blocked with 3% bovine serum albumin in 0.1% Tween 20 PBS and incubated with rabbit anti-Spike antibody (1 in 1000 dilution) for Spike (D614 and G614 variants) detection; with HRP-conjugated StrepTactin (1 in 10000 dilution) for S, E and M detection; or with anti-FLAG (1 in 10000 dilution) for FLAG-tagged S protein. Anti-Spike antibody binding was detected using HRP-conjugated anti-rabbit secondary antibodies (1 in 7000 dilution) (Dako) and visualized using ECL (KPL, Eurobio, Courtaboeuf, France) according to the manufacturer’s instructions. Images were obtained using a G:box imager (Syngene, Fisher Scientific).

### Flow cytometry

HEK cells expressing SARS-CoV-2 S, E and M proteins were resuspended using trypsin (Thermo Fisher Scientific) (1 in 5 dilution in PBS). Cells (2.5×10^5^ per well) were distributed in 96-well plates. For analyzing viral protein expression, cells were fixed and permeabilized following the manufacturer’s instructions (eBiosciences, Thermo Fisher Scientific). HEK cells were then stained with BV650 conjugated streptavidin (1 in 250 dilution) for 30 minutes at 4°C. After washes with a permeabilization buffer, cells were resuspended in PBS 2% FBS and directly analyzed by flow cytometry. For Spike expression, cells were incubated with rabbit anti-Spike antibody for 30 minutes at 4°C, washed three times in PBS 2% FBS, and incubated with AF488 conjugated anti-rabbit antibody for 30 minutes at 4°C. After washes, cells were resuspended in PBS 2% FBS and directly analyzed by flow cytometry. For the serological assay, cells were first incubated with sera (1 in 50 dilution in PBS 2% FBS and 5% donkey serum (PBS FBS/DS)) from healthy donors and SARS-CoV-2 infected patients for 30 minutes at 4°C. Cells were washed in PBS FBS/DS and incubated with AF488 and AF647 conjugated donkey anti-human IgG and IgM F(ab’)2 antibodies or AF488 conjugated goat anti-human IgA F(ab’)2 antibodies for 30 minutes at 4°C. After washing, the cells were resuspended in PBS FBC/DS containing 7AAD reagent (BD Biosciences, Allschwil, Switzerland) to exclude the dead cell population and directly analyzed using flow cytometry on a Novocyte flow cytometer (Acea Biosciences, Agilent). The population of interest was gated according to its FSC/SSC criteria. The dead cell population was excluded using 7AAD staining. Data were analyzed with the NovoExpress software (Acea Biosciences). For protein expression levels, results were expressed as specific fluorescence intensity given by the ratio of the mean of test / the mean of control (*i.e.* secondary antibodies alone). For Ig binding level, results were expressed as specific Ig binding given by the ratio of specific fluorescence intensity obtained with HEK cells expressing viral membrane proteins / specific fluorescence intensity obtained with HEK cells only exposed to the transfection reagent (without DNA).

### Molecular modeling

Sequences used for predicted protein structures of Spike D614 variant (PDB ID 6ZB5, EM 2.85Å resolution) and G614 variant (PDB ID 6XS6, EM 3.70Å resolution, lacking the RBD domain) were initially aligned using ClustalOmega. Sequence alignment showed almost a complete identity except for residue D/G614, an RRA insertion at position 681 in 6BZ5, and a PP→KV mutation at residue 983-984 in 6BZ5. In addition, the initial structural analysis of the G614 variant (6SX6) revealed a clear lack of resolved structures, including the loop between T824 and K851. The homology model (HM) using 6SX6 sequence hence yielded an erroneous geometry. Instead, the structure based on the 6BZ5 sequence with a manually introduced D614G mutation was used. All modeling performed using the Molecular Operating Environment (MOE) 2018.01 software (Chemical Computing Group Inc, Montréal, Canada) and Amber10:EHT force field.

### Statistical analyses

Graphs and statistical analyses were performed using GraphPad Prism 7.0 software (GraphPad Software). Data are presented as mean ± SD or SEM of at least three independent experiments. Statistical significance (p<0.05 or less) was determined using a paired or unpaired t-test or ANOVA when appropriate.

## Results

### Expression of Spike, E and M in mammalian cells and antibody-based detection of mature envelope proteins

As viral membrane protein recognition is part of the anti-SARS-CoV-2 immune response, we developed a mammalian cell-based serological assay using SARS-CoV2 S, E and M expressing human embryonic kidney (HEK) cells to mimic integral membrane protein maturation found at the surface of viral particles (**Figure 1A**). HEK cells were transiently transfected with genes encoding for SARS-CoV-2 S, E and M proteins in tandem with either two Strep-Tag II motifs (11) or a FLAG tag. Forty-eight hours post-transfection, expression of S, E and M was confirmed using both western blotting with HRP-conjugated StrepTactin, anti-FLAG or anti-Spike antibodies, and flow cytometry using BV650-conjugated streptavidin (**Figure 1B**). Cell surface expression of S was also confirmed using flow cytometry using an anti-Spike antibody (**Figure 1C**). The proportion of positive cells and the expression levels of viral proteins were similar between experiments and when viral proteins were compared (**Figure 1C** and **1D**). To validate the binding of anti-SARS-CoV2 IgG, M and A subtypes to HEK cells expressing viral membrane proteins, we used two sera from SARS-CoV-2 infected patients (COVID+ and CTR#4 the latter being distributed by SeroBio as a validation tool for diagnostic laboratories) and one serum from a healthy donor (PRECOV obtained before January 2020) as a negative control. Sera were incubated with non-permeabilized HEK cells and Ig binding was detected using secondary antibodies specific for each Ig subtype. Non-specific binding was determined using non-transfected HEK cells. The detection of IgG, M and A binding was observed on S-expressing HEK cells using sera form SARS-CoV-2 infected patients in a concentration-dependent manner, whereas no Ig binding was found using the healthy serum (**Figures 1E** and **1F**). These results indicate that S is expressed at the surface of HEK cells and can be detected by anti-S antibodies from COVID+ patients.

**Figure 1:**
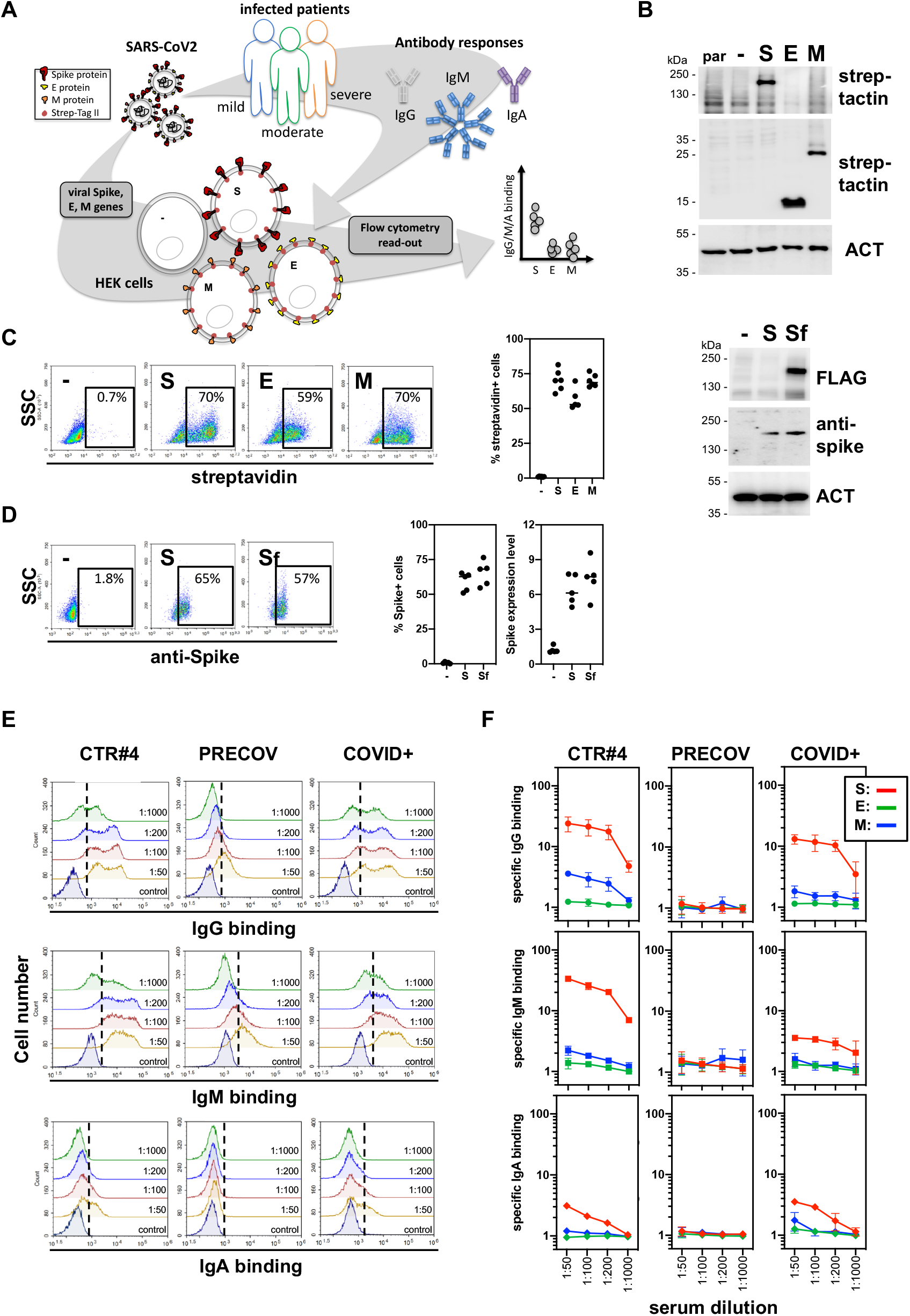
Development of a serological assay that mimics surface expression of SARS-CoV-2 membrane proteins. **(A)** Strategy and workflow of the SARS-CoV-2 serological assay developed using flow cytometry. HEK cells transiently transfected with viral genes encoding membrane proteins (*i.e.* E, M and Spike) were used as matrix for the detection of antibodies present in sera obtained from COVID-19 patients. The binding of anti-viral membrane proteins antibodies of IgG, IgM and IgA subtypes was analyzed by flow cytometry. Results were expressed as specific antibody binding to viral membrane proteins as non-specific binging was determined using non-transfected HEK cells. **(B-D)** Expression of viral membrane proteins at the surface of HEK cells. HEK cells transfected with viral genes encoding E, M and Spike membrane proteins tagged with two Strep Tag motifs were analyzed for viral protein expression by western-blot **(B)** and flow cytometry **(C-D)** using strep-tactin, streptavidin or anti-Spike antibody. Percentage of positive cells and protein expression levels were represented in **(C)** and **(D)**. **(E-F)** Detection of immunoglobulin IgG, IgM and IgA binding at the surface of HEK cells expressing viral membrane proteins by flow cytometry. Positive sera from SARS-CoV-2 infected patients (CTR#4 and COVID+) were incubated with HEK cells expressing E, M and Spike viral proteins at different dilutions. Serum from a healthy donor (PRECOV) obtained before January 2020 was used as a negative control. The binding of IgG, M and A immunoglobulins were analyzed by flow cytometry using secondary antibodies specific for each Ig subtype. Representative flow cytometry histograms were shown for Spike in **(E)** and results were presented as specific Ig binding relative to HEK cells **(F)**.

### Anti-S and M serological responses in COVID+ patients

A cohort of 131 patients was next tested in our serological assay including sera from i) heathy/asymptomatic donors obtained before (n=38) and after January 2020 (n=26), ii) patients infected with non-SARS-CoV-2 coronavirus (n=5), iii) patients suffering from hyper-immunoglobulin M syndromes (n=5), iv) patients with symptoms similar to those observed in COVID+ patients (*i.e.* anosmia, cough, fatigue, fever) (n=4), and v) patients previously infected with SARS-CoV-2 (as confirmed by PCR, n=51) and developing mild (patients who did not need hospitalization, n=22), moderate (hospitalized patients treated with oxygen therapy <5L, n=14) and severe (hospitalized patients in ICU with oxygen therapy >5L or intubated, n=15) forms of the COVID-19 disease (**Table 1**). The time between the PCR tests (or the first symptoms) and the blood sampling were similar between the groups of COVID-19 patients (between 15 and 25 days; **Figure S1A**). Our results were comparable to those obtained with assays developed for diagnostic laboratories by Beckman (IgG anti-Spike) and Roche (Ig anti-N protein), showing 93% and 89% concordance, respectively (**Figure S1B**). No Ig binding to S, E and M proteins was observed in control sera including those obtained before January 2020, those from patients infected with other coronaviruses or suffering from hyper-immunoglobulin M syndromes (**Figures 2A, 2B** and **S1C**). The positive threshold was therefore set using these control sera. Anti-Spike IgG, M and A were detected in sera from COVID+ patients as well as in those from patients with COVID-19-associated symptoms. Anti-Spike Ig titers were higher in patients with moderate and severe forms of the disease compared to mild forms (**Figures 2A** and **2C)**. Anti-E Ig were never detected in any of the tested sera (**Figure S1C**). Importantly, anti-M Ig were also observed at a higher level in patients with severe forms of the disease (**Figures 2B** and **2C)**. Interestingly, while sera positive for anti-M Ig always exhibited anti-Spike Ig signals, some anti-Spike Ig positive sera did not show any detectable anti-M Ig (**Figure2D**).

**Figure 2:**
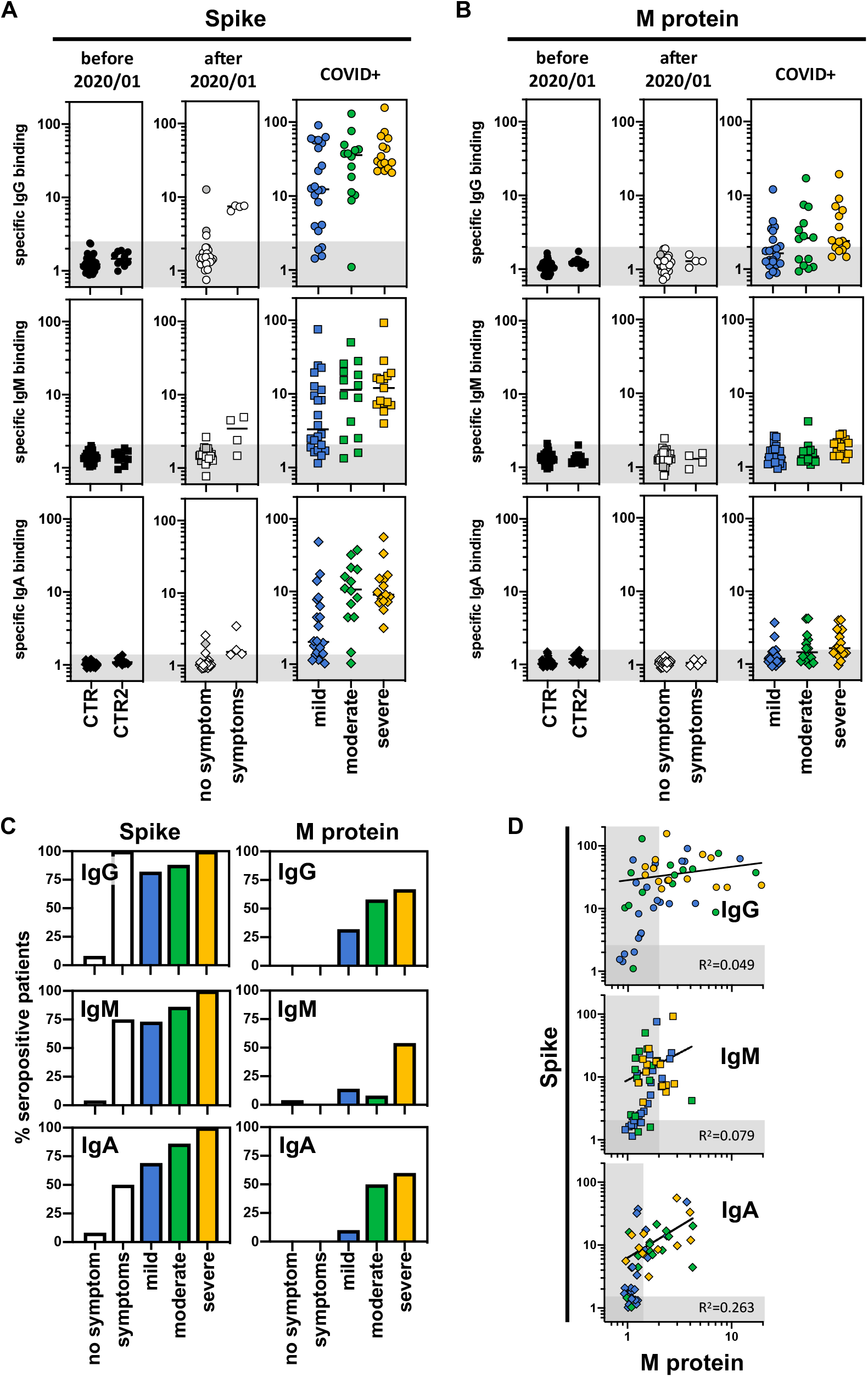
Serological profile of SARS-CoV-2 infected patients against viral membrane proteins according to the disease severity. **(A-B)** Sera from control donors and SARS-CoV-2 infected patients were tested for their positivity against viral Spike **(A)** and M proteins **(B)** using the SARS-CoV-2 serological assay described in Figure 1. Control sera were obtained from healthy donors and collected before January 2020 (CTR, n=38); and from patients infected with other coronaviruses (n=5) or patients with hyperimmunoglobulin M syndrome (n=5) (included in CTR2, n=10). Sera collected after January 2020 were obtained from donors without symptoms (no symptom, n=26); with symptoms related to SARS-CoV-2 infection (symptoms, n=4); and from patients positive for SARS-CoV-2 infection (COVID+) and developing mild (blue, n=22), moderate (green, n=14) and severe (orange, n=15) forms of COVID-19. Specific binding of IgG (circles), IgM (squares) and IgA (diamonds) were represented and thresholds (grey boxes) were obtained with the basal levels of Ig binding from control sera. **(C)** The percentage of seropositive patients from the different groups were calculated using thresholds obtained in **(A-B)**. **(D)** Correlation between anti-Spike and anti-M Ig responses were represented including sera from COVID-19 patients developing mild (blue, n=22), moderate (green, n=14) and severe (orange, n=15) forms of COVID-19.

### Reduced Ig binding to Spike G614 compared to D614 in COVID+ patients

Our assay recapitulating viral protein modification and insertion in a membrane was validated using COVID-19 patients’ sera and yielded results comparable to those obtained with commercially available tests (**Figure S1B**). However, the latter tests use the Spike D614 variant as antigen and it is well established that most European patients until the end of 2020 were mainly infected by SARS-CoV-2 expressing Spike G614 (*e.g.* French patients were exclusively exposed to Spike G614 variant from March to December 2020 (12)). This might lead to biases in data interpretation. Hence, we sought to investigate potential differences in terms of antibody responses using our assay.

First, we compared the structural properties of the Spike D614 and G614 variants. We focused on the beta-sheet rich domain containing D/G614 of chain A (yellow), and its interaction with the patch T824-E865 on chain B (golden), squared in **Figure 3A**. D614 (chain A) forms an inter-protomeric salt bridge to K854 (chain B) (**Figure 3B top**). In the same region, there is also an inter-protomeric salt bridge between R646 (chain A) → E865 (chain B). Using the model with the D614G mutant of 6BZ5, we observed that K854 (chain B) remains pointing towards protomer A, and forms a H-bond to the carbonyl backbone of G614 (*i.e.* a weaker and more strained interaction than in the case of D614 (**Figure 3B, bottom**)). The second salt bridge R646 (chain A) → E865 (chain B) is still retained. Looking at the electrostatic and hydrophobic properties of the area in protomer A, we observed that the domain side interacting with protomer B is largely nonpolar, except for D614 and R646 (**Figure 3C, top**). In contrast, the electrostatic interaction is considerably weaker in the G614 variant, and essentially only retained by R646 protruding towards E854 (chain B) (**Figure 3C, bottom**). The sequences at the domain interfaces consist largely of non-polar residues, and the change from D614 to G614 clearly impacts on the overall polarity of the protomer A interaction area. We also note that in the G614 variant, the loop region after K854 of protomer B (golden) is bending further away from protomer A, than what is observed in the D614 variant. Analysis of the surfaces of protomer B in the interface also illustrates the difference in interactions between the two protomers. In particular the non-polar region of protomer B is protruding towards protomer A between D614 and R646 in the D614 variant, but is in contrast pushed back/out in the G614 variant (**Figure 3D**). In addition the segment around K854 is in G614 clearly rotated away from protomer A. In the region close to E865 of protomer B, an increased exposure of hydrophilic/polar residues towards the solvent (better seen in the lipophilicity surfaces) was observed. These analyses indicate that the G614D mutation might alter the global structure of the protein and therefore the antigenic response.

**Figure 3:**
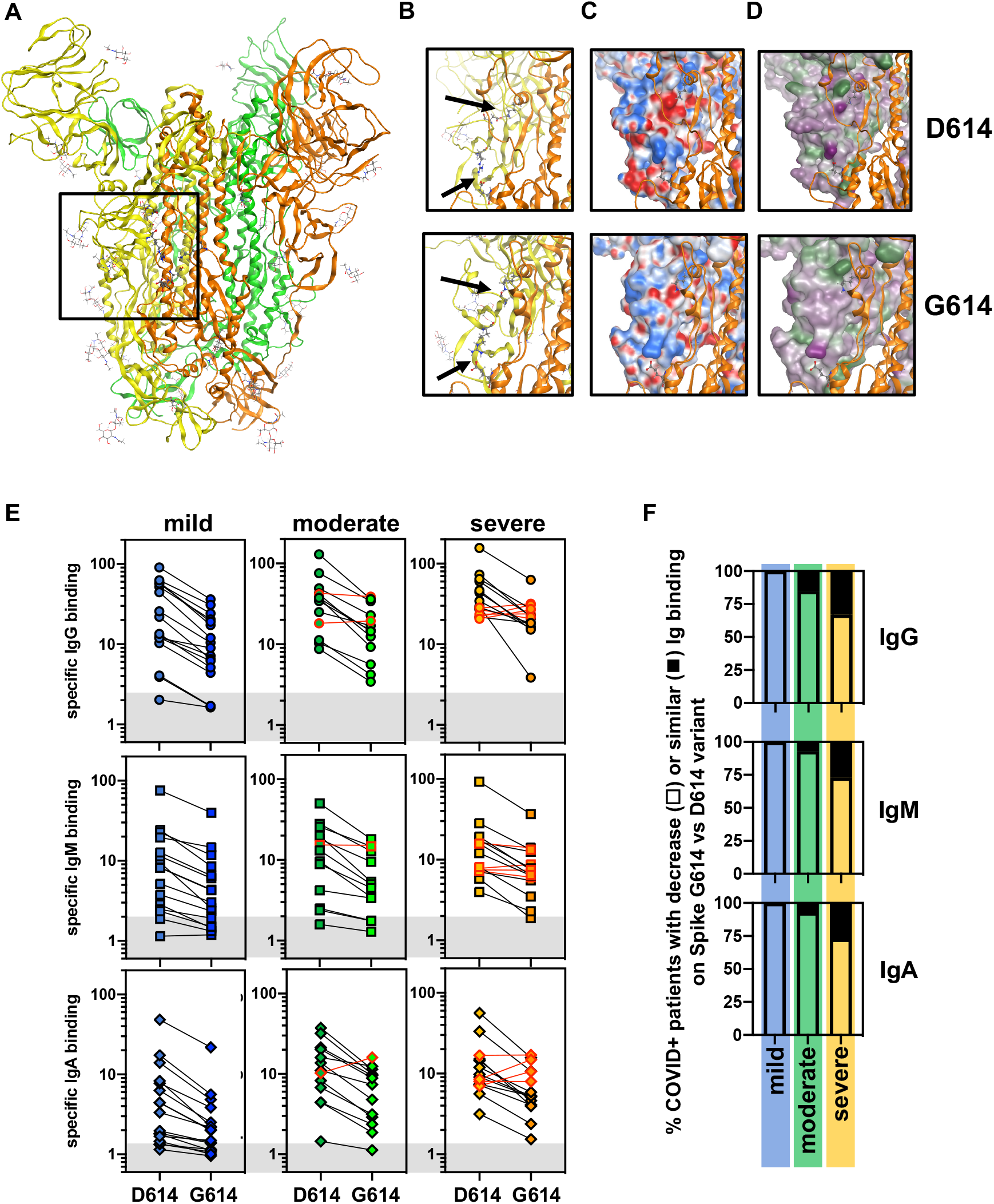
Impact of Spike G614 variant on the seropositivity of SARS-CoV-2 infected patients. **(A-B)** Protein structure of Spike obtained from the Protein Data Bank (PDB ID6zb5 for D614 variant and PDB ID6xs6 for G614 variant). **(A)** The region in which the amino acid 614 was localized (square) on the trimer of Spike molecules (in green, orange and yellow) was further analyzed in D. **(B)** Predicted residue interactions with electrostatic (middle panels) and lipophibic (right panels) properties were compared between Spike D614 and G614 variants. **(C)** Sera from COVID-19 patients developing mild (blue, n=17), moderate (green, n=13) and severe (orange, n=15) forms of COVID-19 were re-evaluated in the SARS-CoV-2 serological assay using Spike D614 and G614 expressing HEK cells. Specific binding of IgG (circles), IgM (squares) and IgA (diamonds) were represented and thresholds (grey boxes) were obtained with the basal levels of Ig binding from control sera tested in Figure 2A. **(D)** Percentage of patients with decreased seropositivity against D614 and G614 variants from the different groups were presented.

Anti-Spike Ig response was therefore re-evaluated using sera from COVID+ patients and anti-Spike D614 and G614 variants expressing cells. Expression of the Spike variants in HEK cells was validated as previously (**Figure S2A-C**). Similar expression levels of Spike D614 and G614 variants were found at the cell surface (**Figure S2C**). Sera from COVID+ patients were then tested on both cell systems and a lower anti-S IgG, M and A binding to Spike G614 was observed in most of the patients than that observed for binding to the D614 variant (**Figure 3E**). Only a small proportion of those patients (less than 15%) displayed similar Ig responses against the two Spike variants (**Figure 3E, F**). Hence our experimental system allows for discrimination between anti-S D614 vs. G614 Ig signals likely due to the advantage of using membrane inserted Spike following complex folding and post-translational modifications.

## Discussion

Using our mammalian cell-based assay with the SARS-CoV-2 envelop proteins S, E and M (**Figure 1**), we identified antibody responses against the S and M proteins in COVID+ patients but not against the membrane viral E protein. Higher anti-S and −M Ig production correlated with symptom severity in hospitalized COVID+ patients (**Figure 2**). Furthermore, whereas French patients were exclusively exposed to SARS-CoV-2 expressing the Spike G614 variant until December 2020, reduced IgG, M and A binding was observed on Spike G614 compared to that observed for the D614 variant (**Figure 3**). Overall, this study underlines the importance of using antigens respecting viral protein constraints to investigate antibody responses against SARS-CoV-2 membrane proteins.

Most of the diagnostic tests currently applied to detect the antibody response against SARS-CoV-2 do target the viral proteins S and N as both were initially found to be expressed abundantly and exhibit substantial antigenicity (13–18). One of the drawbacks associated with these assays is the use of recombinant proteins produced in prokaryote systems that only include fragments of Spike (*e.g.* S1 subunit or RBD domain (13–18)). Recent structural analyses have revealed that the structural integrity of the full-length Spike multimers (trimers, dimers of trimers and more) is important to better understand its immunogenic potential (19). As the virus hijacks the host secretory machinery to produce nascent viral particles in infected cells (3), we designed a reliable serological assay using mammalian cells that express full-length SARS-CoV2 envelope proteins S, M and E, the most exposed to the host immune system. Such a system allows for proper folding and post-translational modifications of the viral proteins (3). These modifications include for instance disulfide bond formation and N-glycosylation thus leading to native structural features.

Recent studies have mapped the regions of SARS-CoV-2 proteins recognized by antibodies from COVID-19 patients using proteome microarrays (20–22). Antibody production was detected against peptides derived from the structural proteins Spike (S1, S2 and RBD domains), N and M and from the accessory protein ORF3a. Interestingly, we are able to demonstrate for the first time the occurrence of an antibody response in COVID+ patients against the entire M protein using our cell-based serological assay, thereby confirming results observed in proteome microarrays in a more physiological context (20–22). Higher Ig binding was observed in hospitalized COVID+ patients with moderate and severe forms of the disease. Of note, Ig response against the M protein was always observed in patients that also exhibited an anti-Spike Ig response. In contrast, no antibody against the E protein was detected in our assay. This could be explained by the fact that the E protein might not be expressed at the surface of the cells or that it may not expose enough antigenic regions to mount a potent immune response. Previous studies indicate that SARS-CoV-2 derived E protein seems to be mainly localized inside the host cells at the ER, ERGIC and/or Golgi compartments although the precise localization is still debated (23, 24).

One additional advantage of the assay developed in this study is the possibility to quickly adapt to express new envelop protein variants. As an example, we compared the Ig responses against the Spike variants D614 and G614. To our surprise, Spike G614 displayed a lower Ig binding capacity compared to D614. Structural analyses revealed differences in electrostatic and hydrophobic surfaces between the Spike variants that could impact on Ig affinity. Interestingly, this mutation is located near the 615 to 635 flexible loop leading to a salt bridge between D/G614 of one protomer and K854 on the neighboring protomer, possibly affecting the global structure of the Spike trimer (25), increasing the RBD ‘up’ state and S1/S2 proteolysis (26). Of note, no difference in Ig binding levels has previously been observed in ELISA-based assays (27). The discrepancy observed with our study could be linked to the type of serological assay used; ELISA versus cell-based assays could exacerbate these protein structural differences. Intriguingly, French COVID-19 patients analyzed in this study were exclusively exposed to SARS-CoV-2 expressing Spike G614 variant, suggesting that the mutation does not reduce the antigenicity against Spike but instead reduce IgG, M and A binding. Many new SARS-CoV-2 strains associated with Spike substitutions have recently emerged (e.g. as observed for variants from Brazil, United-Kingdom and South-Africa) with increased infectivity (28–31). These new strains also display several mutations in other viral genes encoding for structural and accessory proteins. However, very few mutations were described for M so far (32, 33), suggesting that anti-M Ig responses described in this study could be conserved across the different SARS-CoV-2 lineages. If anti-M antibodies effectively reduce SARS-CoV2 infectivity, the low variation burden on this protein might also reveal an efficient tool for vaccine development.

At the current time of massive vaccination against Spike using RNA approach developed by Pfizer-BioNTech and Moderna (34, 35); and the spreading of novel SARS-CoV-2 strains carrying Spike mutations, our serological assay represents a reliable test to verify the immunization efficiency during the vaccination and to analyze the impact of these Spike mutations on antibody responses.

## Acknowledgements

We thank the members of the Virology department of the Rennes University hospital for providing human sera; the Centre Eugène Marquis and INSERM for support. We particularly thank Aurore Nicolas for collecting the clinical data and sera. This work was funded by grants from INSERM, Institut National du Cancer (INCa PLBIO), Fondation pour la Recherche Médicale (FRM, équipe labellisée 2018) to EC and from la Ligue contre le cancer (comité 35, 56 et 85) to TA.

## Author contribution

**SM, GJ**– methodology, investigation, formal analysis; **CH, VT, FG**– resources; **MLG**– conceptualization, writing (review & editing); **LAE**– conceptualization, investigation, formal analysis, modeling of S protein; **EC**– supervision, conceptualization, project administration, funding acquisition; writing (review & editing); **TA**– supervision, conceptualization, methodology, investigation, formal analysis, project administration, writing (original draft, review & editing) (https://www.casrai.org/credit.html#).

## References

1. Hu B, Guo H, Zhou P, Shi ZL. Characteristics of SARS-CoV-2 and COVID-19. Nat Rev Microbiol. 2020.

2. Poland GA, Ovsyannikova IG, Kennedy RB. SARS-CoV-2 immunity: review and applications to phase 3 vaccine candidates. Lancet. 2020;396(10262):1595–606.

3. Sicari D, Chatziioannou A, Koutsandreas T, Sitia R, Chevet E. Role of the early secretory pathway in SARS-CoV-2 infection. The Journal of cell biology. 2020;219(9).

4. Ravi N, Cortade DL, Ng E, Wang SX. Diagnostics for SARS-CoV-2 detection: A comprehensive review of the FDA-EUA COVID-19 testing landscape. Biosens Bioelectron. 2020;165:112454.

5. Zohar T, Alter G. Dissecting antibody-mediated protection against SARS-CoV-2. Nat Rev Immunol. 2020;20(7):392–4.

6. Lan J, Ge J, Yu J, Shan S, Zhou H, Fan S, et al. Structure of the SARS-CoV-2 spike receptor-binding domain bound to the ACE2 receptor. Nature. 2020;581(7807):215–20.

7. Hoffmann M, Kleine-Weber H, Schroeder S, Kruger N, Herrler T, Erichsen S, et al. SARS-CoV-2 Cell Entry Depends on ACE2 and TMPRSS2 and Is Blocked by a Clinically Proven Protease Inhibitor. Cell. 2020;181(2):271–80 e8.

8. Grzelak L, Temmam S, Planchais C, Demeret C, Tondeur L, Huon C, et al. A comparison of four serological assays for detecting anti-SARS-CoV-2 antibodies in human serum samples from different populations. Science translational medicine. 2020;12(559).

9. Ripperger TJ, Uhrlaub JL, Watanabe M, Wong R, Castaneda Y, Pizzato HA, et al. Orthogonal SARS-CoV-2 Serological Assays Enable Surveillance of Low-Prevalence Communities and Reveal Durable Humoral Immunity. Immunity. 2020;53(5):925–33 e4.

10. Yang Q, Hughes TA, Kelkar A, Yu X, Cheng K, Park S, et al. Inhibition of SARS-CoV-2 viral entry upon blocking N- and O-glycan elaboration. Elife. 2020;9.

11. Gordon DE, Jang GM, Bouhaddou M, Xu J, Obernier K, White KM, et al. A SARS-CoV-2 protein interaction map reveals targets for drug repurposing. Nature. 2020;583(7816):459–68.

12. Korber B, Fischer WM, Gnanakaran S, Yoon H, Theiler J, Abfalterer W, et al. Tracking Changes in SARS-CoV-2 Spike: Evidence that D614G Increases Infectivity of the COVID-19 Virus. Cell. 2020;182(4):812–27 e19.

13. Algaissi A, Alfaleh MA, Hala S, Abujamel TS, Alamri SS, Almahboub SA, et al. SARS-CoV-2 S1 and N-based serological assays reveal rapid seroconversion and induction of specific antibody response in COVID-19 patients. Scientific reports. 2020;10(1):16561.

14. Amanat F, Stadlbauer D, Strohmeier S, Nguyen THO, Chromikova V, McMahon M, et al. A serological assay to detect SARS-CoV-2 seroconversion in humans. Nature medicine. 2020;26(7):1033–6.

15. Lee CY-P, Lin RTP, Renia L, Ng LFP. Serological Approaches for COVID-19: Epidemiologic Perspective on Surveillance and Control. Frontiers in immunology. 2020;11(879).

16. Houlihan CF, Beale R. The complexities of SARS-CoV-2 serology. The Lancet Infectious diseases. 2020;20(12):1350–1.

17. Rikhtegaran Tehrani Z, Saadat S, Saleh E, Ouyang X, Constantine N, DeVico AL, et al. Performance of nucleocapsid and spike-based SARS-CoV-2 serologic assays. PloS one. 2020;15(11):e0237828.

18. La Marca A, Capuzzo M, Paglia T, Roli L, Trenti T, Nelson SM. Testing for SARS-CoV-2 (COVID-19): a systematic review and clinical guide to molecular and serological in-vitro diagnostic assays. Reproductive biomedicine online. 2020;41(3):483–99.

19. Bangaru S, Ozorowski G, Turner HL, Antanasijevic A, Huang D, Wang X, et al. Structural analysis of full-length SARS-CoV-2 spike protein from an advanced vaccine candidate. bioRxiv. 2020.

20. Krishnamurthy HK, Jayaraman V, Krishna K, Rajasekaran KE, Wang T, Bei K, et al. Antibody profiling and prevalence in US patients during the SARS-CoV2 pandemic. PloS one. 2020;15(11):e0242655.

21. Wang H, Wu X, Zhang X, Hou X, Liang T, Wang D, et al. SARS-CoV-2 Proteome Microarray for Mapping COVID-19 Antibody Interactions at Amino Acid Resolution. ACS Cent Sci. 2020;6(12):2238–49.

22. Jiang H-w, Li Y, Zhang H-n, Wang W, Men D, Yang X, et al. Global profiling of SARS-CoV-2 specific IgG/ IgM responses of convalescents using a proteome microarray. medRxiv. 2020:2020.03.20.20039495.

23. Satarker S, Nampoothiri M. Structural Proteins in Severe Acute Respiratory Syndrome Coronavirus-2. Arch Med Res. 2020;51(6):482–91.

24. Miserey-Lenkei S, Trajkovic K, D’Ambrosio JM, Patel AJ, Čopič A, Mathur P, et al. A comprehensive library of fluorescent constructs of SARS-CoV-2 proteins and their initial characterization in different cell types. bioRxiv. 2020:2020.12.19.423586.

25. Zhang L, Jackson CB, Mou H, Ojha A, Rangarajan ES, Izard T, et al. The D614G mutation in the SARS-CoV-2 spike protein reduces S1 shedding and increases infectivity. bioRxiv. 2020:2020.06.12.148726.

26. Gobeil SM, Janowska K, McDowell S, Mansouri K, Parks R, Manne K, et al. D614G Mutation Alters SARS-CoV-2 Spike Conformation and Enhances Protease Cleavage at the S1/S2 Junction. Cell reports. 2021;34(2):108630.

27. Klumpp-Thomas C, Kalish H, Hicks J, Mehalko J, Drew M, Memoli MJ, et al. D614G Spike Variant Does Not Alter IgG, IgM, or IgA Spike Seroassay Performance. medRxiv. 2020:2020.07.08.20147371.

28. Leung K, Shum MH, Leung GM, Lam TT, Wu JT. Early transmissibility assessment of the N501Y mutant strains of SARS-CoV-2 in the United Kingdom, October to November 2020. Euro Surveill. 2021;26(1).

29. Tegally H, Wilkinson E, Giovanetti M, Iranzadeh A, Fonseca V, Giandhari J, et al. Emergence and rapid spread of a new severe acute respiratory syndrome-related coronavirus 2 (SARS-CoV-2) lineage with multiple spike mutations in South Africa. medRxiv. 2020:2020.12.21.20248640.

30. Paiva MHS, Guedes DRD, Docena C, Bezerra MF, Dezordi FZ, Machado LC, et al. Multiple Introductions Followed by Ongoing Community Spread of SARS-CoV-2 at One of the Largest Metropolitan Areas of Northeast Brazil. Viruses. 2020;12(12).

31. Gröhs Ferrareze PA, Franceschi VB, de Menezes Mayer A, Caldana GD, Zimerman RA, Thompson CE. E484K as an innovative phylogenetic event for viral evolution: Genomic analysis of the E484K spike mutation in SARS-CoV-2 lineages from Brazil. bioRxiv. 2021:2021.01.27.426895.

32. Mercatelli D, Giorgi FM. Geographic and Genomic Distribution of SARS-CoV-2 Mutations. Frontiers in microbiology. 2020;11:1800.

33. Laamarti M, Alouane T, Kartti S, Chemao-Elfihri MW, Hakmi M, Essabbar A, et al. Large scale genomic analysis of 3067 SARS-CoV-2 genomes reveals a clonal geo-distribution and a rich genetic variations of hotspots mutations. PloS one. 2020;15(11):e0240345.

34. Walsh EE, Frenck RW, Jr., Falsey AR, Kitchin N, Absalon J, Gurtman A, et al. Safety and Immunogenicity of Two RNA-Based Covid-19 Vaccine Candidates. The New England journal of medicine. 2020;383(25):2439–50.

35. Anderson EJ, Rouphael NG, Widge AT, Jackson LA, Roberts PC, Makhene M, et al. Safety and Immunogenicity of SARS-CoV-2 mRNA-1273 Vaccine in Older Adults. The New England journal of medicine. 2020;383(25):2427–38.

