## Supplemental Data for "SARS-CoV2 envelop proteins reshape the serological responses of COVID-19 patients"

**Supplemental Figures legends**

**Figure S1:** *Serological profile of SARS-CoV-2 infected patients against viral membrane proteins according to the disease severity.* **(A)** Times from blood taken and PCR test and symptoms were represented for the different groups of COVID-19 patients used in this study. **(B)** Results obtained with our serological assay was compared to results obtained with serological assays used for diagnosis from Beckman and Roche companies. **(C)** Sera from control donors and SARS-CoV-2 infected patients were tested for their positivity against viral E proteins using the SARS-CoV-2 serological assay described in Figure 1. Control sera were obtained from heathy donors and collected before January 2020 (CTR, n=38); and from patients infected with others coronavirus (n=5) or patients with hyperimmunoglobulin M syndrome (n=5) (included in CTR2, n=10). Sera collected after January 2020 were obtained from donors without symptoms (no symptom, n=26); with symptoms related to SARS-CoV-2 infection (symptoms, n=4); and from patients positive for SARS-CoV-2 infection (COVID+) and developing mild (blue, n=22), moderate (green, n=14) and severe (orange, n=15) forms of COVID-19. Specific binding of IgG (circles), IgM (squares) and IgA (diamonds) were represented and thresholds (grey boxes) were obtained with the basal levels of Ig binding from control sera.

**Figure S2:** *Impact of Spike G614 variant on the seropositivity of SARS-CoV-2 infected patients.* **(A-C)** Expression of viral membrane Spike D614 and G614 proteins at the surface of HEK cells. HEK cells transfected with viral genes encoding Spike D614 and G614 membrane proteins tagged with two Strep Tag motifs were analyzed for viral protein expression by western-blot (A) and flow cytometry (B-C) using strep-tactin, streptavidin or anti-Spike antibody. The percentage of positive cells and protein expression levels were represented in (B) and (C). **(D)** Results obtained with our serological assay with Spike G614 variant was compared to results obtained with serological assays used for diagnosis from Beckman and Roche companies. **(E)** Predictive residue interactions with electrostatic, hydrophobic (left panels) and lipophobic (right panels) properties were compared between Spike D614 and G614 variants.

### FIGURE S1

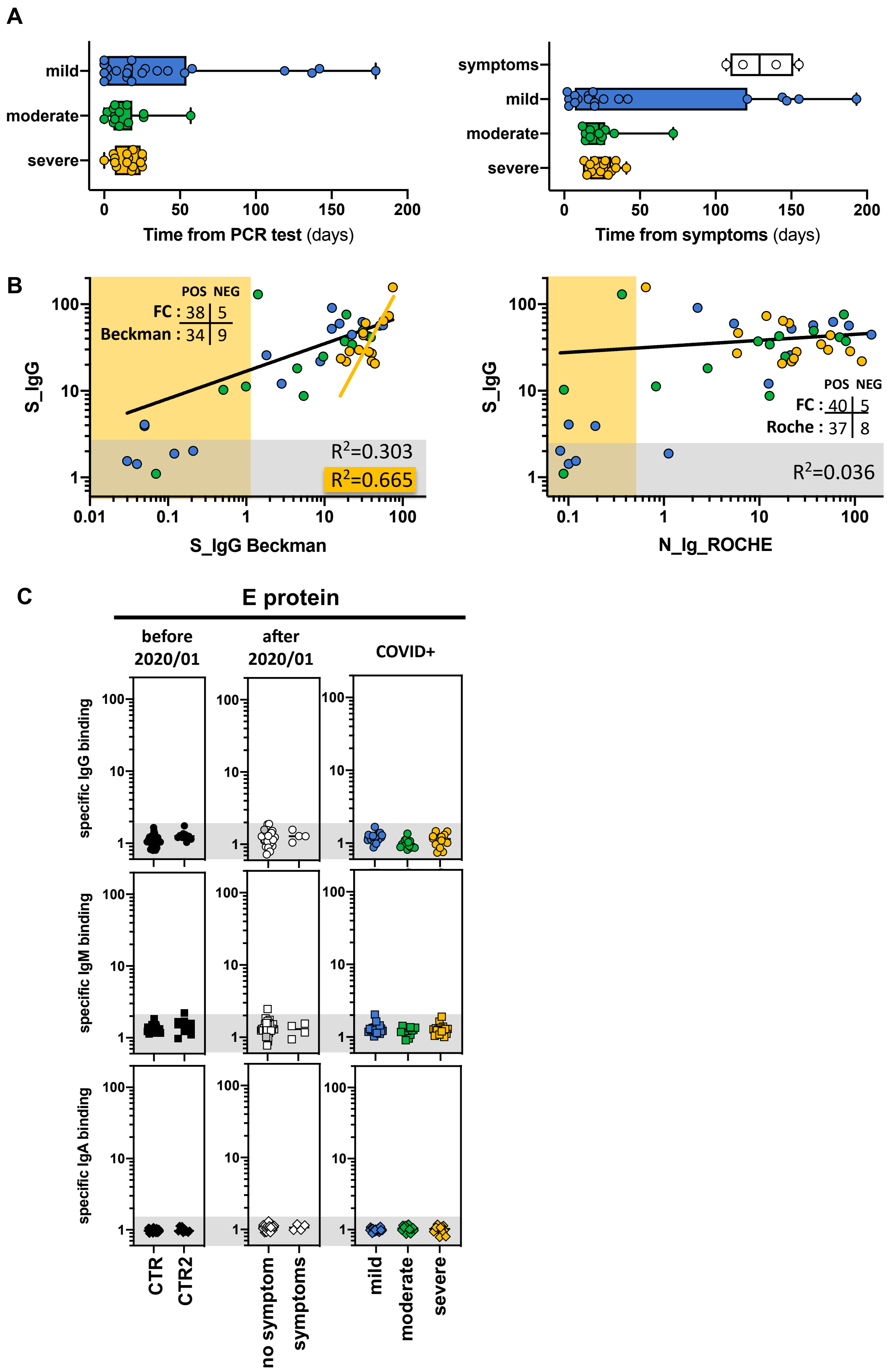

A
mild
moderate
severe
0 50 100 150 200
Time from PCR test (days)
symptoms
mild
moderate
severe
0 50 100 150 200
Time from symptoms (days)
B
S_IgG
POS NEG
FC : 38 | 5
Beckman : 34 | 9
R²=0.303
R²=0.665
0.01 0.1 1 10 100
S_IgG Beckman
S_IgG
POS NEG
FC : 40 | 5
Roche : 37 | 8
R²=0.036
0.1 1 10 100
N_Ig_ROCHE
C
E protein
before 2020/01
after 2020/01
COVID+
specific IgG binding
specific IgM binding
specific IgA binding
CTR
CTR2
no symptom
symptoms
mild
moderate
severe

### FIGURE S2

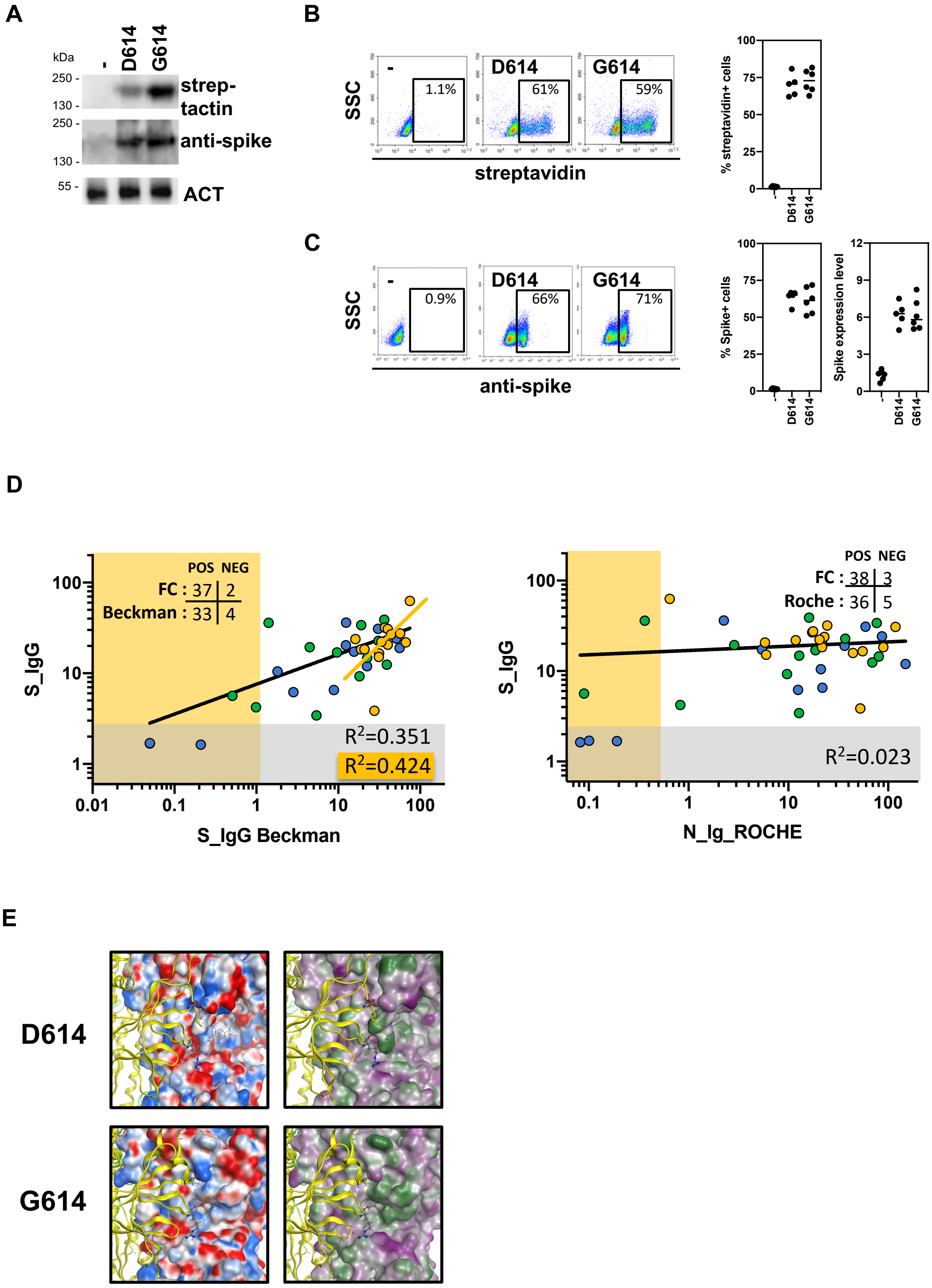
A
kDa
-
D614
G614
250 -
130 -
250 -
130 -
55 -
strep-
tactin
anti-spike
ACT
B
SSC
-
1.1%
D614
61%
G614
59%
streptavidin
% streptavidin+ cells
C
SSC
0.9%
66%
71%
anti-spike
% Spike+ cells
Spike expression level
D
POS NEG
FC : 37 | 2
Beckman : 33 | 4
S_IgG
R2=0.351
R2=0.424
S_IgG Beckman
POS NEG
FC : 38 | 3
Roche : 36 | 5
R2=0.023
N_Ig_ROCHE
E
D614
G614
